# Heterogeneity of Sox2-expressing cells in mouse pituitary and their roles in postnatal gonadotroph differentiation

**DOI:** 10.1101/2025.06.03.657631

**Authors:** Kosara Smiljanic, Stephanie Constantin, Naseratun Nessa, Stanko S. Stojilkovic

## Abstract

Postnatal differentiation of gonadotrophs from Sox2-expressing stem cells is essential for maturation of the hypothalamic-pituitary-gonadal axis, puberty, and reproduction. Here, we examined the differentiation and maintenance of gonadotrophs in developing and adult female mice. Gonadotrophs and Sox2-expressing cells were visualized by immunostaining, and gonadotrophs were also identified by specific expression of the fluorescent protein tdTomato during embryonic and postnatal development. Sox2-expressing cells are localized in the anterior parenchyma, marginal zone, and posterior pituitary, regardless of mouse age. Gonadotrophs are localized in the anterior parenchyma distinct from Sox2-expressing cells. During the juvenile and prepubertal periods, cells in transition from Sox2 expression to the tdTomato expression, as well as numerous differentiated gonadotrophs, were also present in the marginal zone. The size and distribution of newly differentiated gonadotrophs was consistent with their maturation and migration into parenchyma. Specific knockout of PI4 kinase A in gonadotrophs disrupted their postnatal differentiation in the marginal zone, causing a significant reduction in the size of the gonadotroph population. This was accompanied by a progressive loss of gonadotroph-specific gene/protein expression and an increase in the number of regressed tdTomato-expressing cells. Thus, cells expressing Sox2 in the marginal zone, but not in the parenchyma, serve as stem cells for postnatal gonadotrophs, and the differentiation and maintenance of these cells require PI4 kinase A-derived phosphoinositides.

## 2. Introduction

The role of stem/progenitor cells (hereafter referred to as stem cells) in the development of the embryonic pituitary is well established (Perez Millan et al., 2024, Alatzoglou et al., 2020). In 2005, initial evidence was also provided that the adult pituitary contains a putative stem cell population (Chen et al., 2005). A series of studies supporting this hypothesis followed (Fauquier et al., 2008, Gleiberman et al., 2008, Garcia-Lavandeira et al., 2009, Chen et al., 2009). Both in vitro and in vivo studies have shown that SRY-box transcription factor 2 (Sox2) expressing cells function as pituitary stem cells during the embryonic and postnatal periods (Fauquier et al., 2008, Andoniadou et al., 2013, Rizzoti et al., 2013). Morphological and transcriptomic heterogeneity of postnatal Sox2-positive cells has been previously suggested for the rodent pituitary (Yoshida et al., 2016a). Sox2-expressing cells line the marginal zone, called marginal cell layer (MCL), but also appear in the parenchyma of the anterior lobe (Gremeaux et al., 2012, Chen et al., 2013). We have also reported that Sox2 gene and protein are expressed in the rat pituicytes of the posterior pituitary (Fletcher et al., 2023).

The main difference between embryonic and postnatal pituitary stem cells is related to the expression of the S100b gene and protein (Soji et al., 1994). It was initially reported that rat folliculostellate cells (FSCs) in the parenchyma express this protein postnatally (Nakajima et al., 1980). This was followed by the generation of S100β/GFP-transgenic rats (Itakura et al., 2007), further characterization of FSCs (Horiguchi et al., 2010), and their ability to differentiate into hormone producing cells in vitro (Higuchi et al., 2014). It has recently been reported that Sox2-expressing cells in both MCL and parenchyma express *S100b* (Horiguchi et al., 2021), which supports the function of both Sox2 cell populations as stem cells (Vankelecom, 2010). Several single cell RNA sequencing (scRNAseq) studies of the mouse pituitary have also been interpreted in favor of all Sox2-expressing anterior pituitary cells as stem cells (Cheung et al., 2018, Lopez et al., 2021, Vennekens et al., 2021, Sheridan et al., 2025). These findings have questioned the heterogeneity of Sox2-epressing pituitary cells, and the existence of FSCs and their specialized functions typical of differentiated cells (Le Tissier and Mollard, 2021). It is also unknown which pituitary cell types differentiate postnatally from stem cells, only gonadotrophs or other cell types, and which subpopulation of Sox2-expressing cells, parenchymal or MCL, is responsible for their differentiation (Sheridan et al., 2025, Fu et al., 2012).

Here we studied the heterogeneity of Sox2-expressing cells in mouse pituitary tissue and their roles in postnatal differentiation of gonadotrophs. This anterior pituitary cell lineage is specialized for synthesis and release of follicle-stimulating hormone (FSH) and luteinizing hormone (LH), which are essential for sexual development and reproduction (McArdle and Roberson, 2015). These hormones are heterodimers consisting of a common alpha chain and hormone-specific beta chains (FSHB and LHB), encoded by the *Cga*, *Fshb*, and *Lhb*, respectively (Cahoreau et al., 2015). Gonadotrophs are also defined by the expression of gonadotropin-releasing hormone (GnRH) receptor gene (*Gnrhr*) (Hapgood et al., 2005, Janjic et al., 2017). This receptor is activated by the decapeptide GnRH, which is synthetized by a subpopulation of hypothalamic neurons and released into the hypothalamic-pituitary portal system (Herbison, 2018). The activated receptor signals through Gq/11-phospholipase C pathway, which leads to the breakdown of phosphatidylinositol 4,5-bisphosphate (PI(4,5)P2) to inositol-1,4,5-trisphosphate (InsP3) and diacylglycerol (Naor and Huhtaniemi, 2013). This signaling pathway ultimately controls the expression of *Fshb*, *Lhb*, and *Gnrhr*, as well as the synthesis and release of gonadotropins determined by the pattern of endogenous GnRH release or its external application (Constantin et al., 2022).

To examine the heterogeneity of Sox2-positive pituitary cells and their roles in postnatal gonadotroph differentiation, we performed immunostaining analysis of intact pituitary tissue from juvenile, prepubertal, and young and mature adult mice for the expression of Sox2, LHB, and FSHB. Gonadotrophs were also visualized by specific expression of the fluorescent protein tdTomato via GnRH receptor-driven Cre/Lox recombination of the Rosa-tdTomato locus. To investigate the potential role of plasma membrane phosphoinositide content in postnatal gonadotroph differentiation and function, we also used transgenic mice with a gonadotroph-specific phosphatidylinositol 4-kinase alpha (PI4KA) knockout (GSKO), as previously described (Constantin et al., 2023). The numbers of gonadotrophs during the postnatal period were estimated by counting tdTomato-positive cells and qRT-PCR analysis of tdTomato gene expression. We also characterized the time course of postnatal expression of the gonadotroph-specific genes *Fshb*, *Lhb*, and G*nrhr* in control and GSKO mice. The results of these studies indicate that marginal cells serve as stem cells for gonadotrophs and that their postnatal differentiation and function depend on PI4KA-derived phosphoinositides.

## 2. Material and methods

### 2.1. Animals

Pi4ka^fl/fl^ mice (MGI:6294260) (Bojjireddy et al., 2014) were bred with Rosa^tdTomato/tdTomato^ mice (MGI:3809524) (Madisen et al., 2010) to generate double homozygous Pi4ka^fl/fl^ Rosa^tdTomato/tdTomato^ bucks. Pi4ka^fl/fl^ mice were also bred with Gnrhr^IRES-Cre/IRES-Cre^ mice (MGI:3795249) (Wen et al., 2008) to generate Pi4ka^fl/wt^ Gnrhr^IRES-Cre/IRES-Cre^ dams. The transgenic mice generated by their breeding were conditional knockout in which PI4KA is inactivated in GnRH receptor-expressing cells, including gonadotrophs, i.e. Pi4k^fl/fl^ Rosa^tdTomato/wt^ Gnrhr^IRES-Cre/wt^ and their control littermates PI4k^fl/wt^ Rosa^tdTomato/wt^ Gnrhr^IRES-Cre/wt.^. Regardless the transgenic status, the reporter tdTomato was present in cells that express, or have at any time expressed, the *Gnrhr*-driven Cre recombinase. The faithfulness of the Cre reporter in pituitary was assessed in control littermates. Mice were genotyped using the KAPA HotStart mouse genotyping kit (Kapa Biosystems, Sigma-Aldrich, St. Louis MO) and 0.5 μM specific primers, as described previously (Constantin et al., 2023). Mice were housed under constant temperature and humidity, with a standard 14-hour light/10-hour dark cycle. The sex and age of animals used for experiments is stated in figures and/or figure legends. All experiments with animals described in this study were performed in accordance with the National Institutes of Health Policy Manual 3040-2: Animal Care and Use in the Intramural Program and were approved by the National Institute of Child Health and Human Development, Animal Care and Use Committee (Animal Protocol 22-041).

### 2.2. qRT-PCR analysis

Pituitaries were collected from mice at different time points: postnatal day (week) 11, 14, 18, 21, 42 (6), 70 (10), 98 (14), 154 (22), 182 (26) to detect gene expression changes over time. Each pituitary was processed individually, except at postnatal day 11 for which each datapoint required 2 pituitaries. Mice were euthanized by an overdose of isoflurane (Baxter, Deerfield, IL), the pituitary glands were collected and snap frozen on dry ice. Total RNA from pituitaries were isolated with RNeasy Plus Mini Kit (Qiagen, Valencia, CA), reverse transcribed with Transcriptor First Strand cDNA Synthesis Kit (Roche Applied Sciences, Indianapolis, IN), and qRT-PCR was performed using Applied Biosystems pre-designed TaqMan Gene Expression Assays (Applied Biosystems, Waltham, MA), according to manufacturer’s instructions. Target gene expression levels were determined by the comparative 2^-(delta Ct) quantification method using *Gapdh* as the reference gene (Janjic et al., 2019). Applied Biosystems predesigned TaqMan Gene Expression Assays were used: *Fshb* (Mm00433361_m1), *Gapdh* (Mm99999915_g1), *Gnrhr* (Mm00439143_m1), *Lhb* (Mm01205505_g1), and *tdTomato* (Mr07319439).

### 2.3. Immunostaining

Mice were anesthetized with ketamine/xylazine cocktail (200/20 mg/kg body weight), intracardially perfused with 0.1 M PBS, then with PBS containing 4% formaldehyde. Pituitaries were collected, postfixed overnight, and transferred to PBS containing 30% sucrose until the tissue sank. Pituitaries were cut on a Leica CM3050 S cryostat, 12 µm sections were mounted on SuperFrost slides (Fisher Scientific, Waltham, MA) and stored on -80°C. On the day of experiment, pituitary sections were warmed at room temperature for 2 h. After washing 2 times in PBS, antigen retrieval was performed. Slides were incubated in Universal HIER antigen retrieval reagent (Abcam, ab208572) for 30 min on 80°C, followed by cooling on room temperature for 30 min. All primary antibodies were applied overnight at 4°C, and appropriate secondary antibody for 1 h on RT. Antibodies used in this study are following: rabbit anti-FSHB antibody (RRID:AB_2687903) and guinea pig anti-LHB antibody (RRID:AB_2665565) (both obtained from Dr. A. F. Parlow, National Institute of Diabetes and Digestive and Kidney Diseases, National Hormone and Peptide Program, Torrance, CA; 1:250), rabbit anti SOX2 antibody (RRID:AB_2038021; 1:100), Alexa Fluor 488 goat anti-rabbit (RRID:AB_143165; 1:1000), Alexa Fluor 568 goat anti-guinea pig (RRID:AB_2534119; 1:1000), and Cy5 goat anti guinea pig (RRID:AB_10710629; 1:1000). All antibodies diluted in 1xPBS containing 0.5% BSA and 0.5% Triton X-100. Every step is followed with 4×5 min washing in 1xPBS. Nuclear staining was done with 4′,6-diamidino-2-phenylindole (DAPI) from Invitrogen (REF.00-4959-52). Images were acquired using a Zeiss LSM 900 with Airyscan 2 confocal microscope (Carl Zeiss GmbH, Jena, Germany). For quantification, fluorescence images were analyzed with IMARIS (Oxford Instruments, Abingdon, UK). Images were submitted to auto-thresholding to avoid user’s bias. The total number of cells was estimated by surface detection on the DAPI channel and automatic counting. The number of LHB- and tdTomato-positive cells was estimated by surface detection in 488 nm and 555 nm channels, respectively. The number of cells co-labeled with LHB and tdTomato was estimated by selecting LHB surfaces near the tdTomato surface (<1 µm). The proportion of LHB, tdTomato, or LHB/tdTomato cells was expressed as a percentage of the total number of cells.

### 2.4. Statistical analysis

Results are presented as representative histological images or mean ± SEM values from at least three similar experiments, and the number of replicates were stated in Results or Figure Legends. Statistical analysis was performed by Student *t*-tests for comparison between age-matched control and GSKO groups, and one-way ANOVA for data and post hoc Šidák’s multiple comparison test for data containing more than two groups, with at least P < 0.01 considered statistically significant. Initial graphs were generated using the KaleidaGraph program (Synergy Software, Reading, PA) and figures were finalized using Adobe Illustrator and Photoshop (Adobe, San Jose, CA).

## 3. Results

### 3.1. Characterization of gonadotroph population

The focus of our study is on the postnatal differentiation of gonadotrophs in the mouse anterior pituitary. To identify these cells in a mixed population of pituitary cells, we used two experimental approaches. First, anterior pituitary sections were immunostained using specific LHB and FSHB antibodies as described in Methods. LHB was well detected in pituitary sections from young adult female mice, and LHB-positive cells represented approximately 8% of all anterior pituitary cells (Fig. 1A). The FSHB antibody used in our study also recognized gonadotrophs (Fig. 1B). Combined immunostainings of LHB and FSHB clearly showed that both antibodies recognized the same cells, although the intensity of FSHB staining was variable across gonadotrophs (Fig. 1C). Since both antibodies labeled gonadotroph cytoplasm and our FSHB antibody did not provide additional information about gonadotroph subpopulations, we chose the LHB antibody for further work in this study.

**Fig. 1.**
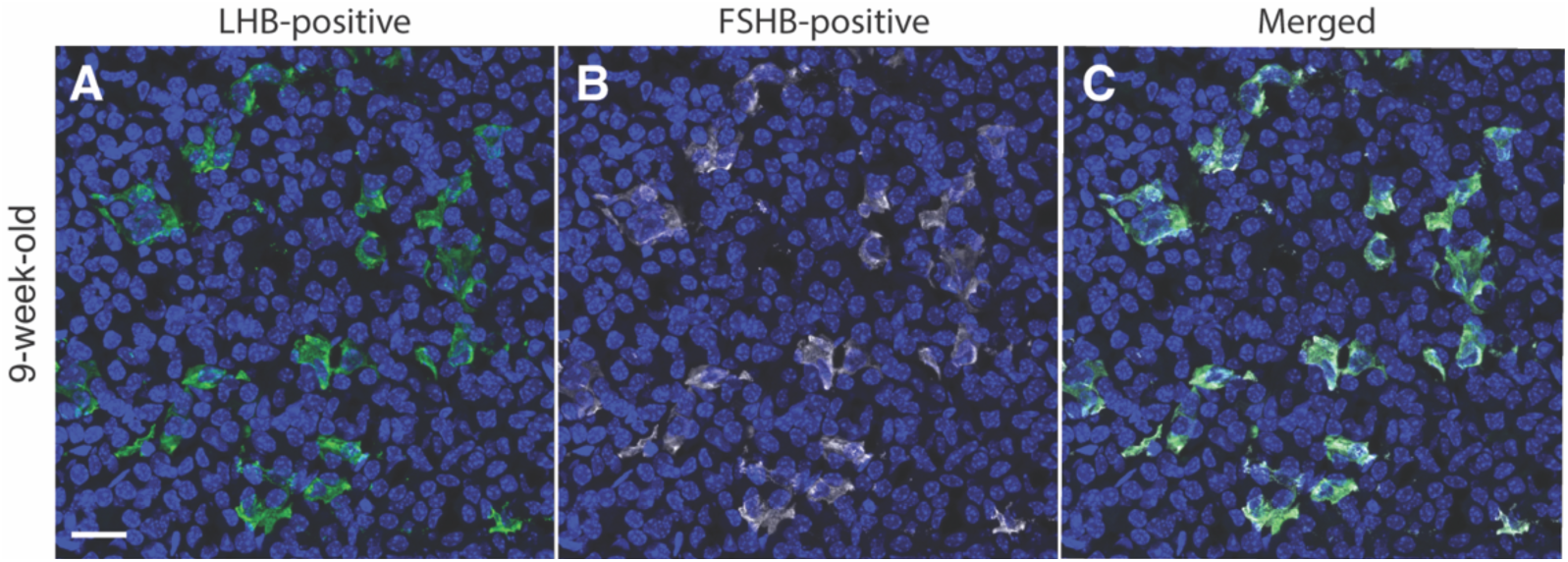
Identification of gonadotrophs in anterior pituitary tissue from young adult female mice. Pituitary gonadotrophs were identified by endogenous expression of LHB (A, green) and FSHB (B, silver) determined by immunostaining. Blue indicates nuclei of all cells labeled by Dapi. Merged of two panels shows that identical cells were labeled (C). For details see Material and Methods. The scale bar of 20 μm shown in panel A applies to all panels.

Second, all our experiments were performed with mice expressing the tdTomato reporter in gonadotrophs via GnRHR-driven Cre/Lox recombination of the Rosa-tdTomato locus, as previously described (Constantin et al., 2023). The expressed tdTomato is a red fluorescent protein commonly used in cell and tissue imaging due to its high brightness and photostability (Shaner et al., 2004). We compared LHB and tdTomato labeling in anterior pituitary sections obtained from prepubertal, young adult, and mature adult animals using female mice aged four (Fig. 2A-C), nine (Fig. 2D-F), and 22 weeks (Fig. 2G-I), respectively. The left panels show LHB immunostaining, the middle panels show tdTomato fluorescent cells, and the right panels show their merging. All LHB-positive cells in the three age groups were also tdTomato-positive, confirming the validity of the tdTomato reporter in identifying gonadotrophs during postnatal mouse life. We also observed the presence of approximately 2% LHB-negative but tdTomato-positive cells (indicated by arrows in Fig. 2), which likely reflects LHB expression below the threshold for detection by immunostaining or minimal ectopic Cre expression. Because the Rosa gene is ubiquitously expressed, Rosa-driven tdTomato expression persists until cell death. Furthermore, tdTomato labeled gonadotroph nuclei in combination with LHB labeled cytoplasm provide double labeling for gonadotroph identification. (Fig. 1E). In this study, some images show only LHB staining or tdTomato fluorescence for simplicity of presentation.

**Fig. 2.**
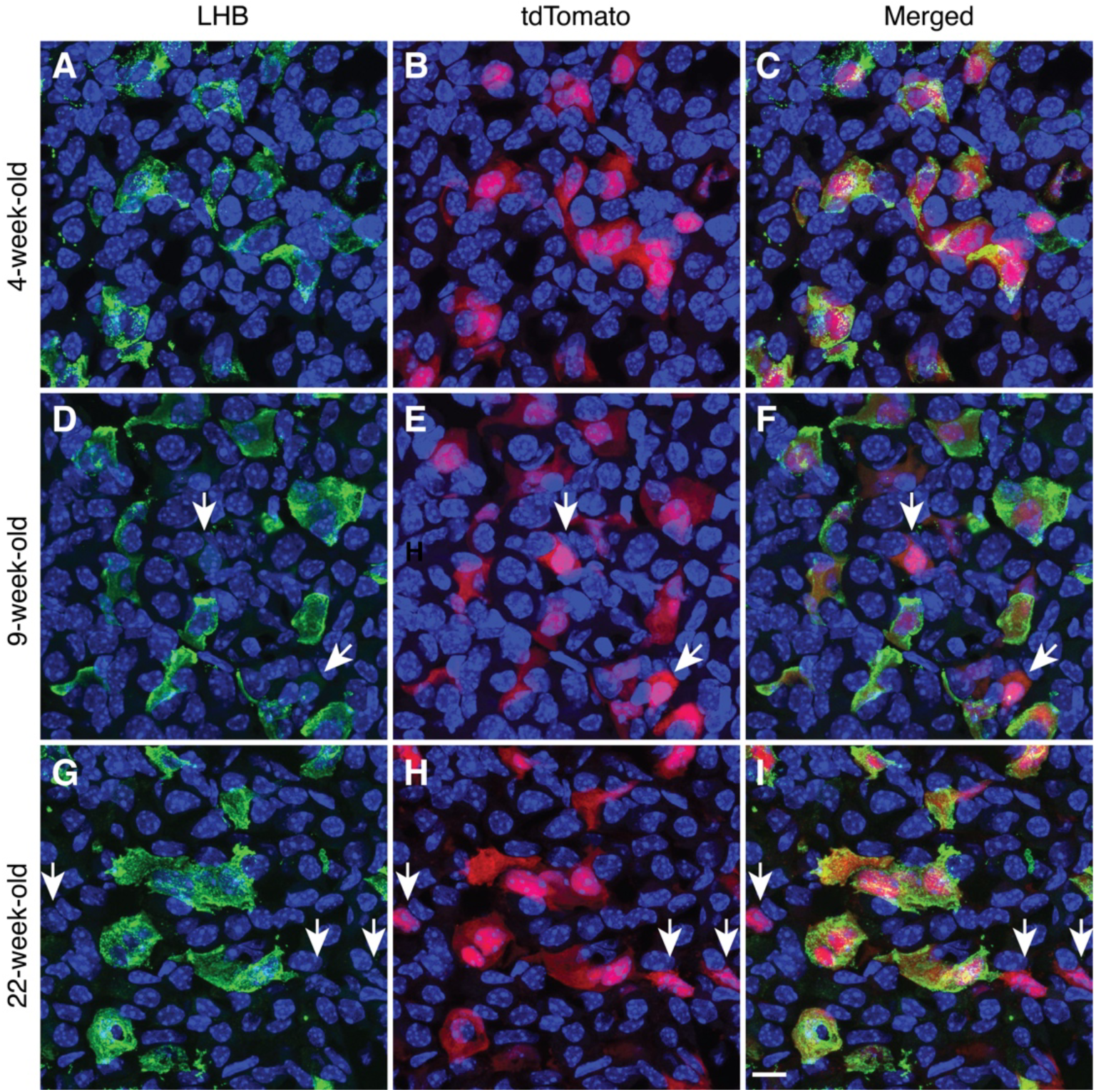
Characterization of tdTomato as an age-independent indicator of postnatal gonadotrophs. Expression of LHB (left panels), tdTomato (middle panels), and their merge (right panels) in anterior pituitary tissues from prepubertal (A-C), young adult (D-F), and mature adult female mice (G-I). Two types of gonadotrophs were observed: cells expressing LHB+ tdTomato and tdTomato-positive cells in which LHB was below the level of detection (indicated by arrows). The 10 μm scale bar shown in panel I refers to all panels.

### 3.2. Postnatal differentiation of gonadotrophs from marginal cells

The rodent pituitary gland consists of three lobes: the anterior lobe, the intermediate lobe, and the posterior lobe. The anterior and intermediate lobes are separated by the marginal zone, which is a remnant of Rathke’s pouch, and contains a layer of cells lining the pituitary cleft (Yoshida et al., 2016a). Figure 3A illustrates the three lobes and the marginal zone in pituitary tissue from a young adult female (9-week-old). Sox2-positive cells (magenta) were detected in the anterior lobe, marginal zone, and posterior lobe, showing a different distribution pattern in these regions. In the anterior lobe, Sox2-expressing cells were randomly distributed as single cells or small cluster of cells. In contrast, Sox2-positive marginal cells were compressed and formed a stream-like band. In the posterior lobe, the majority of pituicytes were Sox2-positive. Taken together, the number of Sox2-expressing cells relative to the total number of cells (233 out of 1888) in this image represented approximately 12%.

**Fig. 3.**
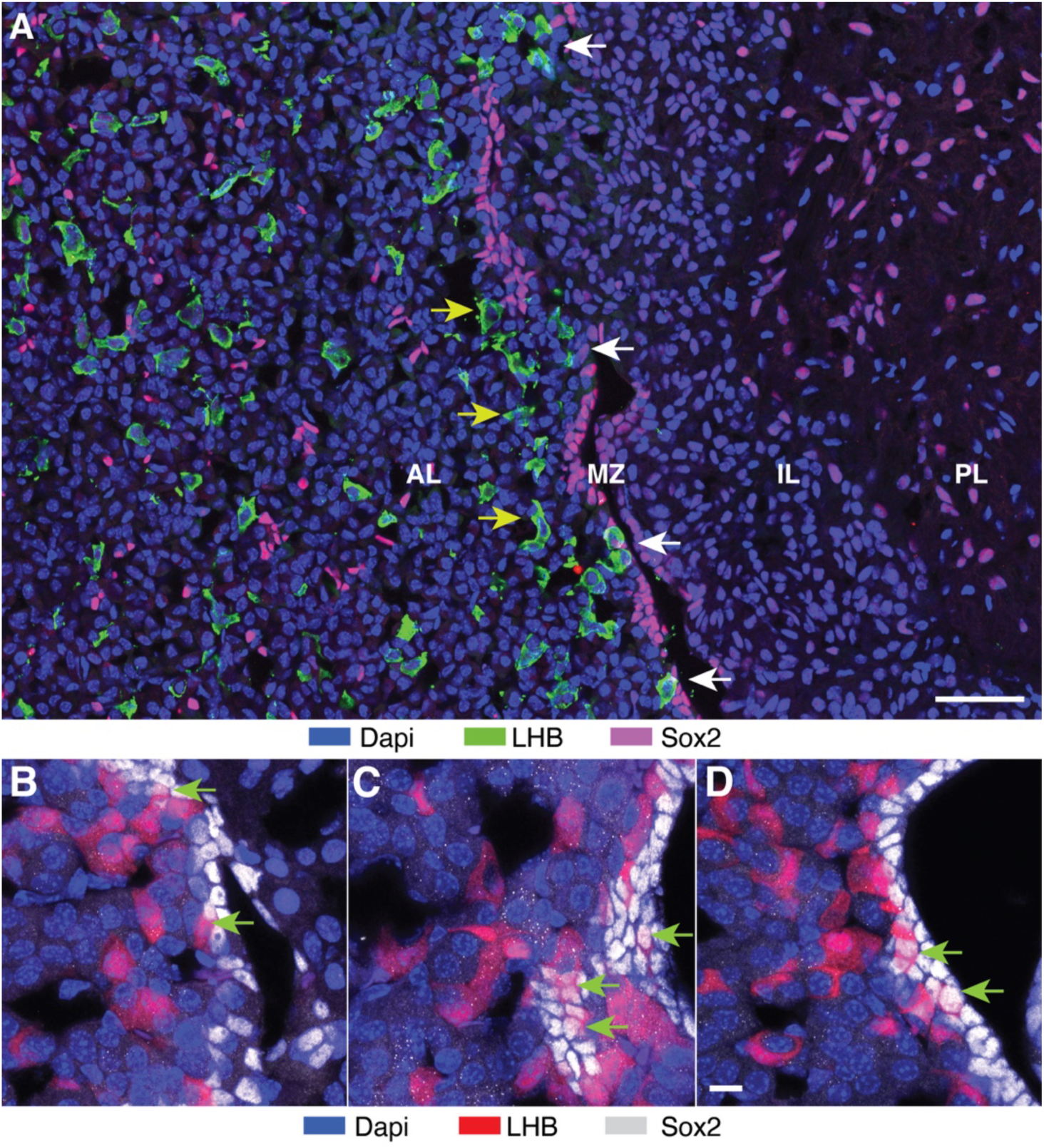
Identification of stem cells accounting for postnatal gonadotroph differentiation. **(A)** Distribution of Sox2-positive cells in the mouse pituitary. Immunostaining of the pituitary gland from 9-week-old female mice: anterior lobe (AL), marginal zone (MZ), intermediate lobe (IL) and posterior lobe (PL). Sox2-positive cells were localized in the AL, MZ, and PL; LHB-positive cells (green) were localized in the AL separately from Sox2-positive cells and in the MZ, some overlapping with Sox2-positive cells, indicated by white arrows. Gonadotrophs in te vicinity of marginal cells, indicated by yellow arrows. The scale bar = 50 µm. (B-D) Transition from Sox2-expressing cells to gonadotrophs in the MZ. Immunostaining of the pituitary of 4-week-old female mice. Green arrows indicate cells in transition, with Sox2 still visible in the nucleus and tdTomato appearing in the cytoplasm. The 10 μm scale bar shown in panel D also applies to B and C panels.

Since gonadotrophs differentiate from Sox2-positive cells (Sheridan et al., 2025), these two cell types should be in close proximity to each other in the marginal zone and/or parenchyma of the anterior lobe. Figure 3A compares the distribution of gonadotrophs (green) and Sox2-positive cells (magenta) in these regions of the anterior pituitary gland. In the parenchyma, both cell types were randomly distributed, but there was no clear pattern of relationship between them. In contrast, some gonadotrophs were mixed with Sox2-expressing cells in the marginal zone, as indicated by the white arrows in Fig. 3A In addition, numerous gonadotrophs were organized as a band parallel to the MCL (pointed by the yellow arrows), suggesting their early step in migration from marginal zone.

Figures 3B-D shows an enlarged marginal zone in pituitary tissue from a prepubertal female (4-week-old), with Sox2 expressing cells shown in silver and tdTomato-labeled cells shown in red. Numerous overlaps of Sox2-expressing cells and gonadotrophs were present in the marginal zone. Green arrows indicate cells in transition from Sox2-expressing to gonadotrophs, as illustrated by the appearance of tdTomato expression and the still visible Sox2 expression in the nucleus. Such a change was not observed in the parenchyma. These data indicate that Sox2-expressing marginal cells serve as postnatal stem cells for pituitary gonadotroph differentiation.

We also performed immunostaining analysis of pituitary tissues from juvenile and prepubertal females, when the gonadotroph population is rapidly increasing, and from mature adult animals, when gonadotroph differentiation has virtually ceased (Sheridan et al., 2025). Figure 4 shows immunostaining of Sox2-positive cells (magenta) and LHB-positive gonadotrophs (green) in the marginal zone and surrounding parenchyma from juvenile (18-day-old), prepubertal (4-week-old) and mature adult (22-week-old) female mice. In pituitary sections from juvenile and prepubertal mice, marginal cells and gonadotrophs overlapped frequently and gonadotrophs were also present in parenchyma close to marginal zone (Fig. 4A and B). However, in mature adult female mice, there were no LHB-positive cells in the MCL and adjacent parenchyma (Fig. 4C). Such an organization is consistent with the conclusion that gonadotrophs differentiate from marginal Sox2-expressing cells during development, emerge from the marginal zone, and gradually migrate deeper into the parenchyma.

**Fig. 4.**
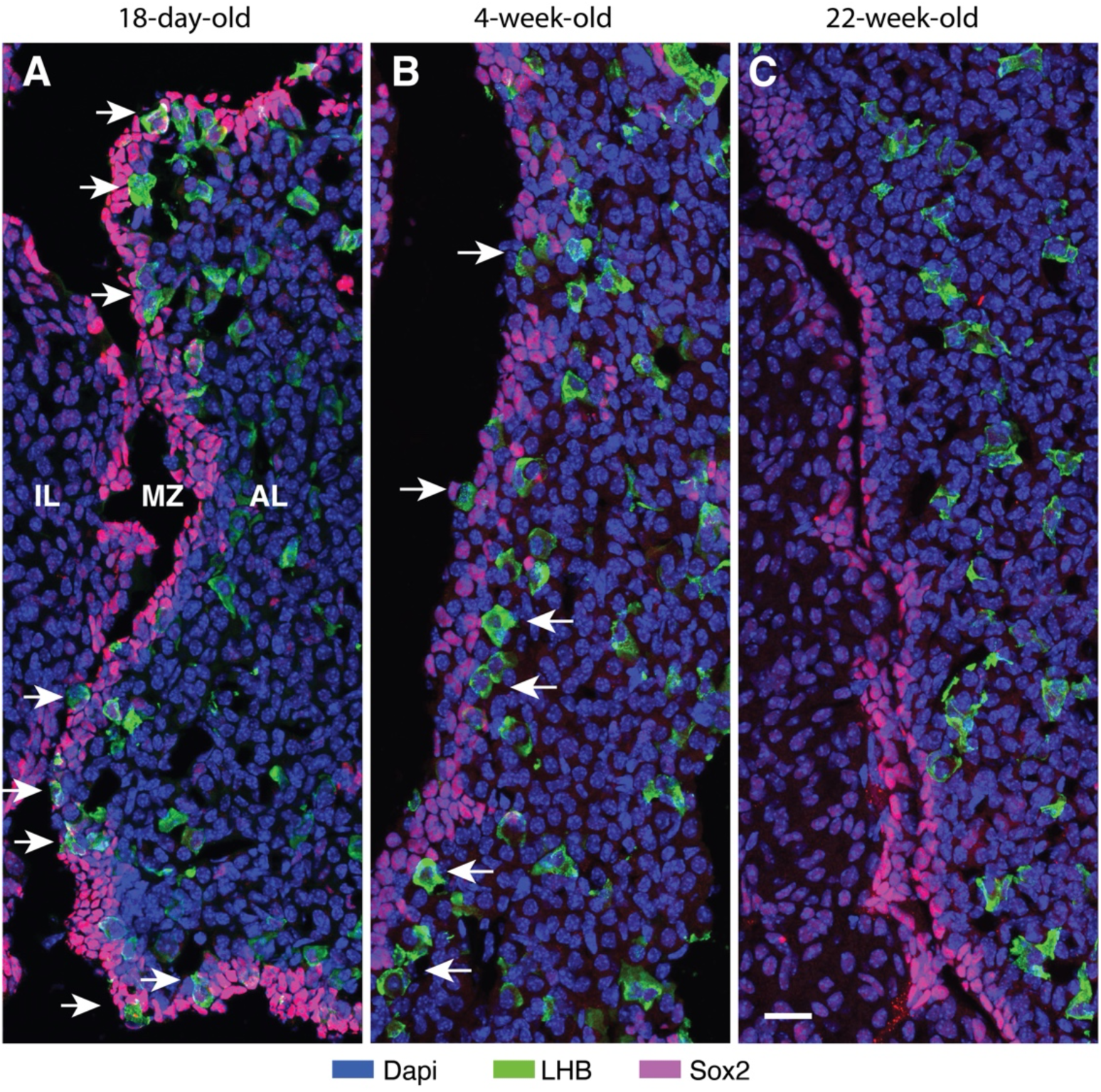
Age-dependent presence of gonadotrophs in the marginal cell layer of female mice. (A - C) Juvenile (18-days, A), prepubertal (4-weeks, B). and mature adult (22-weeks, C) pituitary. IL, intermediate lobe, MZ, marginal zone; AL, anterior lobe. Arrows indicate the presence of LHB in or near the marginal cell layer. Note the difference in diameter of gonadotrophs in the marginal cell layer and the anterior parenchyma. The 20 μm scale bar shown in panel C applies to all panels.

### 3.3. PI4KA dependence of postnatal gonadotroph differentiation

To elucidate the potential role of PI4KA in postnatal differentiation of gonadotrophs from Sox2-expressing stem cells and the size of the gonadotroph population in GSKO mice, we performed immunostaining of control and GSKO pituitary tissues from female mice of different postnatal ages. Figure 5 shows well-developed MCL from three age groups of control (A-C) and GSKO mice (D-F), indicating that the number and organization of stem cells were not affected by GSKO and the age of the animals. However, the presence of transitional cells and differentiated gonadotrophs in the MCL was reduced in prepubertal GSKO pituitaries (Fig. 5A and D). There was also a trend towards a decrease in overlap of gonadotrophs with marginal cells in GSKO pituitaries from 9-week-old females (Fig. 5B vs. E) E). In both controls (Fig. 5C) and GSKO pituitaries (Fig. 5F) of 22-week-old females, there were no cells in transition and differentiated gonadotrophs in the MCL and surrounding parenchyma. In all age groups, the total number of gonadotrophs in GSKO pituitaries was lower compared to age-matched controls. These data show that postnatal gonadotroph differentiation is impaired by PI4KA knockout, suggesting the importance of phosphoinositides generated by this enzyme in postnatal differentiation of gonadotrophs.

**Fig. 5.**
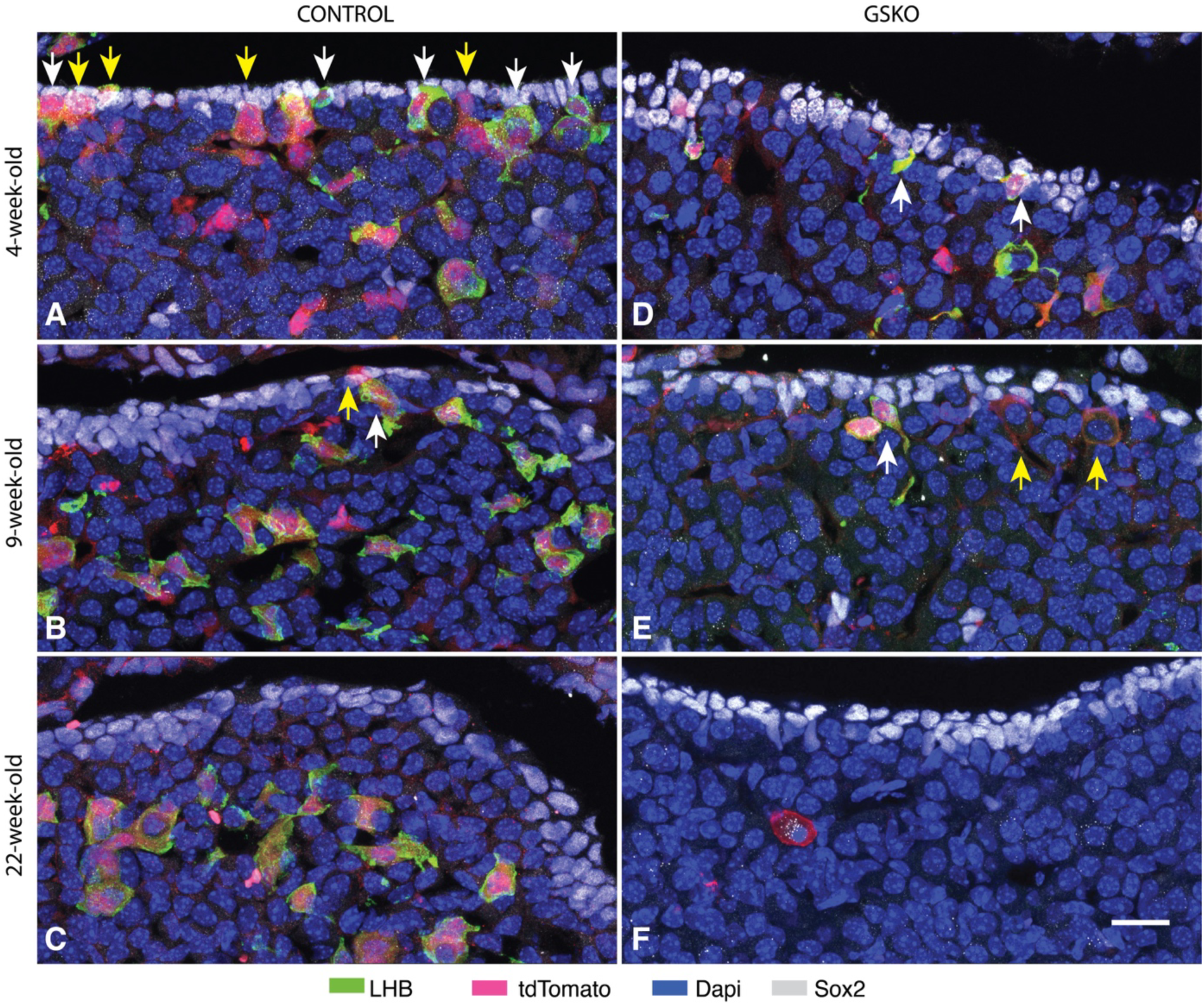
Effects of GSKO on gonadotrophs in the marginal zone. Representative images of gonadotrophs in control (A-C) and GSKO pituitaries (D-F) from juvenile (A and D), young adult (B and E), and mature adult female mice (C and F). Sox2-positive cell layers are well formed in all control and GSKO pituitaries, whereas tdTomato-positive cells (yellow arrows) and tdTomato+LHB-positive cells (white arrows) were detected only in tissue from younger mice (A, B, D, and E). Note the reduction in the number of tdTomato-positive cells overlapping with marginal cells in GSKO pituitaries. The 20 μm scale bar shown in F applies to all panels.

Figure 6 compares gonadotroph populations in the control and GSKO anterior parenchyma at the same age groups. In both control and GSKO pituitary, the number of LHB-expressing gonadotrophs was highest in prepubertal females and decreased in pituitaries from adult animals. However, in all age groups of GSKO females, the number of LHB-expressing gonadotrophs was lower than in control females. Furthermore, the number of cells positive for tdTomato alone remained relatively constant with age in control mice but appeared to increase with age in GSKO mice (indicated by arrows in Fig. 6).

**Fig. 6.**
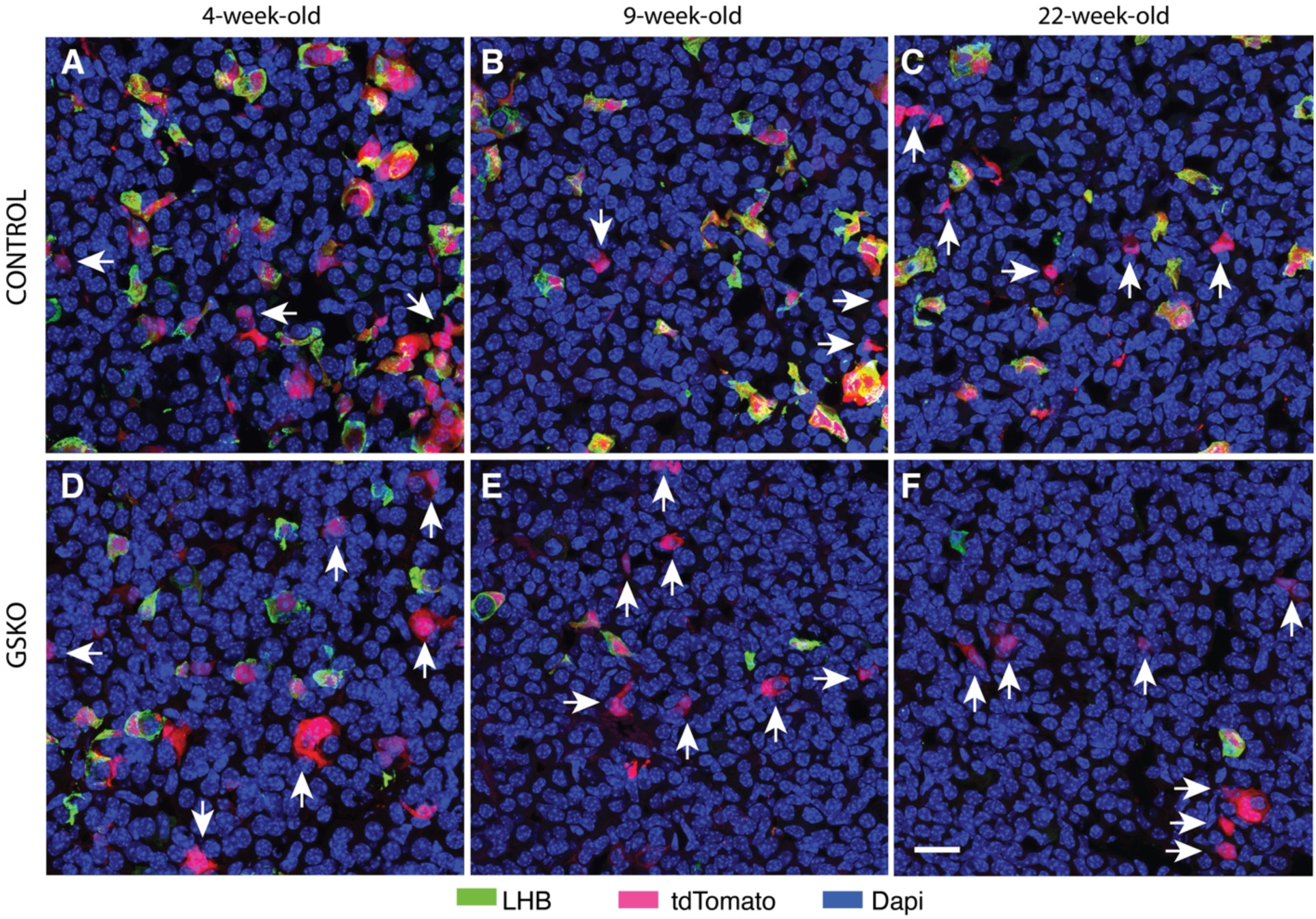
Effect of GSKO on the gonadotroph population in the anterior parenchyma. Representative images of gonadotrophs in control (top panels) and GSKO pituitaries (bottom panels) from prepubertal (left panels) young adult (middle panels), and mature adult female mice (right panels). Gonadotrophs were identified by expression of LHB and/or tdTomato. Arrows indicate only tdTomato positive cells. The 20 μm scale bar shown in F applies to all panels.

To statistically address the issue of the size of the gonadotroph populations in the controls and GSKO pituitaries, we calculated the number of cells positive for LHB + tdTomato relative to cells positive for tdTomato alone and expressed them as a percentage of the total number of cells. These data are shown in Fig. 7 as means ± SEM values. In all age groups, the number of cells positive for LHB + tdTomato was significantly lower in GSKO pituitaries. In contrast, the number of cells positive for tdTomato alone was significantly higher in GSKO pituitaries.

**Fig. 7.**
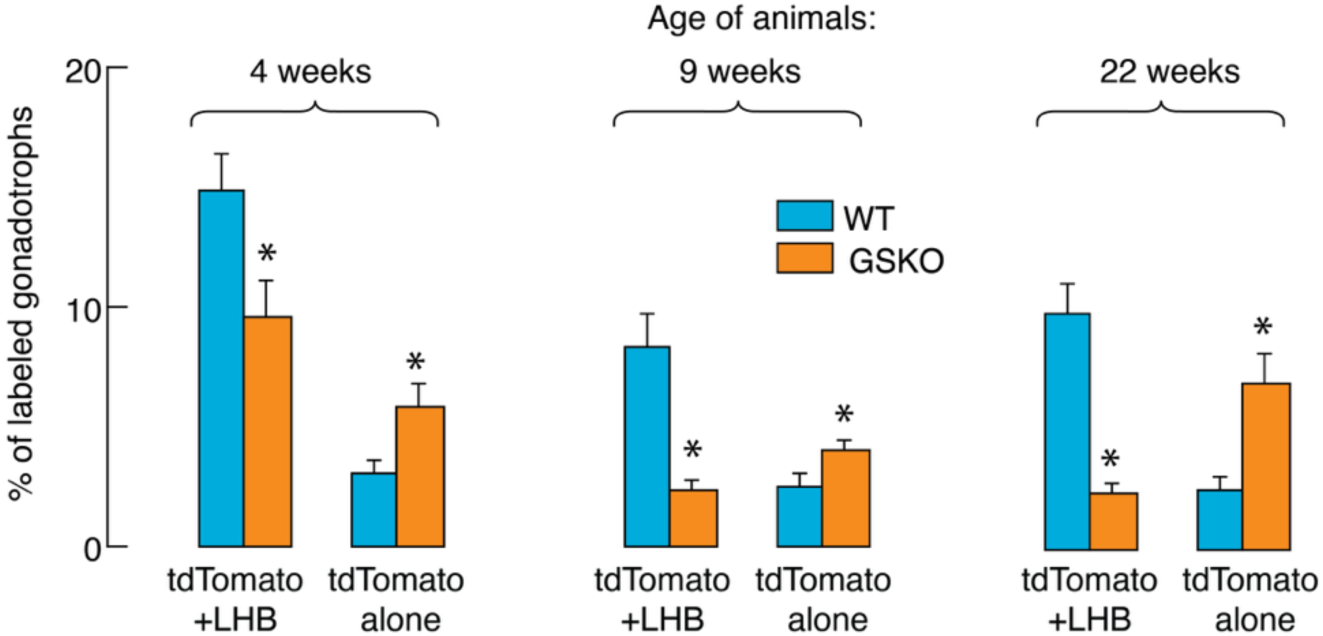
Quantitative characterization of gonadotroph populations in control and GSKO female mice during the postnatal period. Comparison of the percentage of tdTomato+LHB and tdTomato alone expressing gonadotrophs in the anterior pituitary of prepubertal (left), young adult (middle panel) and mature adult (right panel) female mice. Data shown are mean ± SEM, P < 0.01.

We also examined postnatal expression of *tdTomato,* which was used as a reporter for gonadotroph population, in controls and GSKO pituitaries from both females and males (Fig. 8A and B). The temporal profile of *tdTomato* expression was similar in pituitaries of both sexes, with approximately 50% inhibition of expression of this gene in GSKO pituitaries. The squares in panel A illustrate a similar percentage reduction in all tdTomato-positive cells in control and GSKO pituitaries, suggesting that *tdTomato* expression accurately reflects the number of gonadotrophs in pituitary tissue. Thus, GSKO progressively reduced the size of the postnatal gonadotroph population compared with age-matched control animals, reflecting a reduction in postnatal gonadotroph differentiation.

**Fig. 8.**
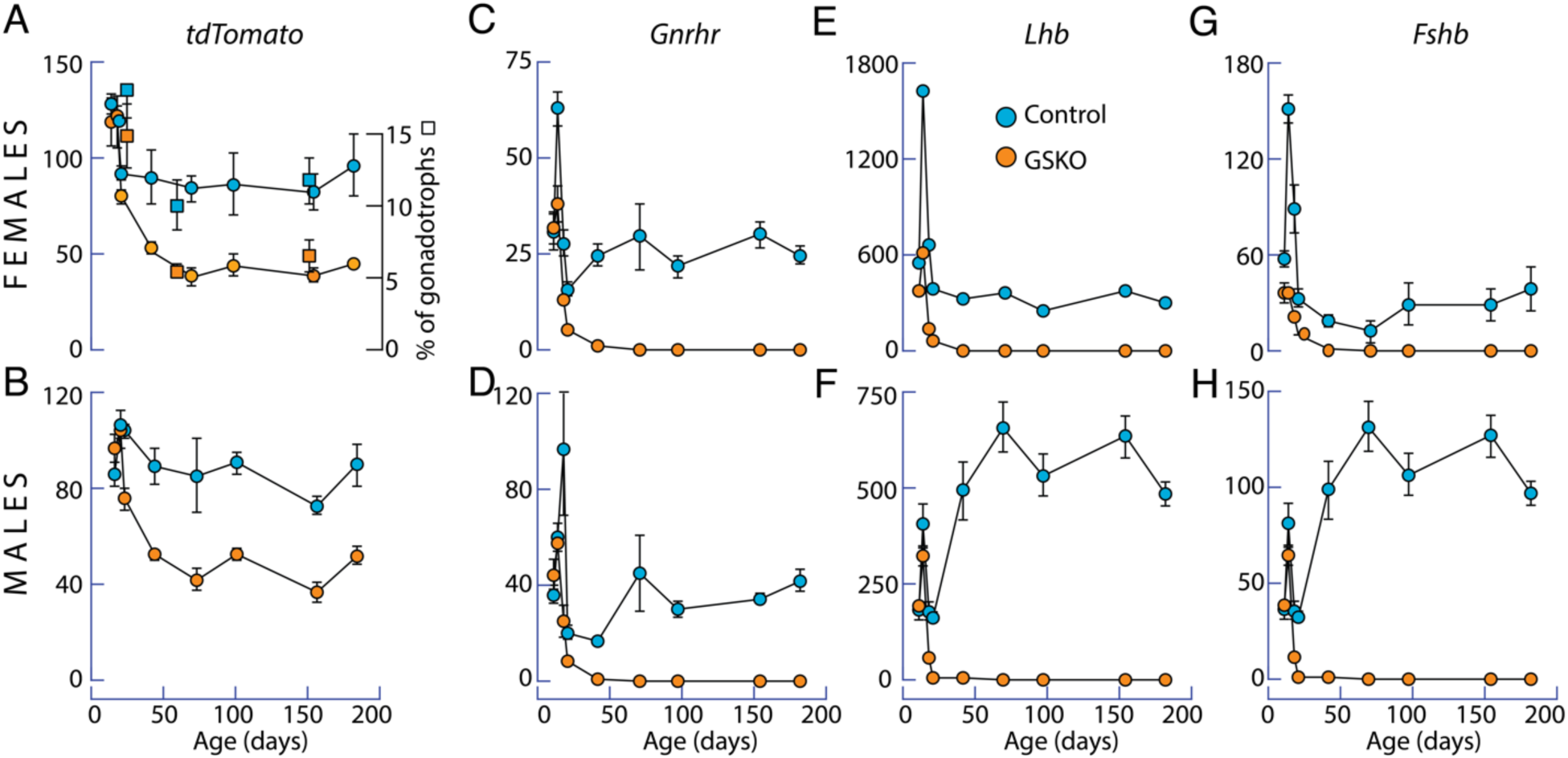
Developmental expression pattern of gonadotroph-specific genes in vivo in control and GSKO mice. (A and B) Similar expression profile of *tdTomato* in male and female pituitaries. GSKO reduced *tdTomato* expression by approximately 50%. Squares illustrate similar percentage reduction in tdTomato-positive cells. (C and D) Comparable expression pattern of *Gnrhr* in female and male controls. (E and F vs. G and H). Sex-specific expression pattern of *Lhb* and *Fshb* in controls. In both sexes, expression of these genes was completely abolished within the first 50 days of postnatal life in GSKO pituitaries. qRT-PCR analysis of gene expression was performed on pituitaries showing tdTomato fluorescence in gonadotrophs and the results are expressed relative to *Gapdh* expression. The number of total tdTomato-positive cells was cells was derived from histological preparations. Data shown are mean ± SEM, with N = 4 - 10 per time point.

Finally, we examined the expression status of gonadotroph-specific genes. The temporal profiles of *Gnrhr* expression in female and male controls were comparable, consisting of an early spike phase and a sustained plateau phase of lower amplitude (Fig. 8C and D). However, the expression of *Lhb* and *Fshb* in control groups was sex specific. In females the profile was comparable to *Gnrhr* expression, and in males, the sustained plateau phase was of higher amplitude than the initial spike phase (Fig. 8E and F vs. G and H). In GSKO pituitaries of both sexes, the expression of these genes was completely abolished within the first 50 days of postnatal life. Thus, GSKO not only reduces the size of the postnatal gonadotroph population compared to age-matched control animals but also blocks the expression of gonadotroph-specific genes in the residual cells.

## 4. Discussion

The identification of Sox2-expressing pituitary cells has been useful in characterizing their location, gene expression, and function during postnatal development of the rodent pituitary (Yoshida et al., 2016a, Davis et al., 2013). It is generally accepted that postnatal Sox2-expressing cells are in the MCL and parenchyma of the anterior pituitary (Gremeaux et al., 2012, Chen et al., 2013). Our previous work with rat pituitary is consistent with these findings but also indicated Sox2 expression in the pituicytes of the posterior pituitary (Fletcher et al., 2023). Here, we show that Sox2-expressing cells are detected in the mouse MCL, anterior parenchyma, and posterior pituitary pituicytes. The distribution of Sox2-expressing cells in the mouse pituitary was similar during juvenile and prepubertal development, as well as in young and mature adult mice. The location and estimated percentage of cells expressing Sox2 relative to the total number of cells in the analyzed sections are consistent with their heterogeneity in terms of expression and function of common and specific genes/proteins.

For example, the S100b gene and protein expression is a common feature of Sox2-positive cells in the anterior and posterior pituitary (Fletcher et al., 2023). Several additional genes were also identified as common to anterior and posterior Sox2-expressing cells in rats, including the development and differentiation genes *Sox9, Cd9*, *Hes1*, *Ntrk2* and *Six3,* and the astrocyte marker genes *Cldn10, Gpr37l1, Fam107a, Fstl1, Glul*, *Sdc4*, and *Slc1a3* (Fletcher et al., 2023). However, anterior and posterior *Sox2*-positive cells in rats also show apparent transcriptomic heterogeneity; *Prop1*, *Prrx1*, *Pitx1* and *Pitx2* are expressed in the anterior but not the posterior pituitary, and *Fgf10*, *Lhx2*, *Rax*, and *Tbx* are expressed in pituicytes but not in *Sox2*-positive anterior pituitary cells (Fletcher et al., 2023). Therefore, the co-expression of most genes in *Sox2*-positive anterior and posterior pituitary cells and their comparable reprogramming is not inconsistent with their separate embryonic development and postnatal differentiation and function.

In contrast, it has been proposed that cells expressing Sox2 in both anterior lobe locations are stem cells, termed the marginal zone niche and the parenchymal niche of stem cells or “primary” and “secondary” niches, respectively (Vankelecom, 2010). Consistent with this hypothesis, several scRNAseq studies of whole or anterior mouse pituitaries have identified Sox2-positive cells as a single cell group, rather than as separate stem cells and FSCs (Cheung et al., 2018, Lopez et al., 2021, Vennekens et al., 2021, Sheridan et al., 2025, De Vriendt et al., 2025). Also, a small fraction of mouse Sox2-expressing cells from parenchyma have been shown to proliferate in vitro under specific experimental conditions (Fu et al., 2012, Yoshida et al., 2016b). It is also well established that the postnatal pituitary contains stem cells that express Sox2 (Perez Millan et al., 2024), and it has recently been shown that gonadotrophs develop postnatally from Sox2-expressing cells (Sheridan et al., 2025).

If postnatal stem cell proliferation leads to differentiation of hormone-producing cells, we hypothesized that cells in transition from stem cells to endocrine cells and newly differentiated endocrine cells should be in the vicinity of Sox2-positive cells in the marginal niche and/or parenchymal niche. To address this question, we chose gonadotrophs, which were visualized by the expression of tdTomato. Like other pituitary endocrine cell types, gonadotrophs also initially emerge in the embryonic pituitary, with early appearance of a common alpha chain and late appearance of FSHB and LHB chains (Zhu et al., 2007, Wen et al., 2010). Postnatally, there is also a progressive growth of gonadotroph population from birth to puberty, with cells spreading dorsally throughout the gland (Mollard et al., 2012, Bjelobaba et al., 2015)..

Because gonadotroph population growth is rapid during the juvenile to peripubertal period and slows down in adult mice, we selected appropriate ages of animals for these experiments. Here we presented evidence of cells switching from Sox2 expression to tdTomato/LHB expression and frequent overlap of Sox2- and LHB-positive cells in the MCL of juvenile and prepubertal females. A shift from Sox2-expressing cells to gonadotrophs, with prominent Sox2-positive nuclei and tdTomato-positive cytoplasm were also evident in a proportion of cells in the MCL. The overlap progressively decreased postpubertally and virtually disappears in mature adult mice. These data indicate that postnatal gonadotrophs originate from Sox2-expressing cells in MCL and that their differentiation occurs in marginal zone before their migration in parenchyma. In contrast, no transition from Sox2-expressing cells to differentiated gonadotrophs, no overlap of Sox2-expressing cells and gonadotrophs in parenchyma of developing and adult mice, nor was there any accumulation of gonadotrophs in specific parenchymal area, arguing against their roles in postnatal gonadotroph differentiation.

The process of postnatal gonadotroph differentiation resembles the neonatal migration of Sox2/S100b-expressing cells from the MCL to the parenchyma, which was originally considered as stem cell migration (Yoshida et al., 2016a). The physiological significance of this process for postnatal animals is unclear. First, the gonadotroph population shows a marked increase during development, and our data clearly indicate that their differentiation in the marginal zone is sufficient to complete this process. Second, the cell turnover rate in the adult pituitary is very low, approximately 1.6% per day (Nolan et al., 1998, Levy, 2002), which is consistent with our observation of a low number of differentiating gonadotrophs seen in adult mice pituitary. Third, ignoring the existence of FSCs does not solve the problem of the well-established specific role of these cells in pituitary physiology (Le Tissier and Mollard, 2021). In accordance with this view, the presence of two postnatal populations of Sox2-positive cells in the anterior pituitary was interpreted as stem cells associated with MCL and differentiated FSCs located in the parenchyma (Fauquier et al., 2008).

Taken together, it is difficult to distinguish between FSCs and stem cells. S100b is expressed in the anterior pituitary after birth (Soji et al., 1994) and was originally considered a marker gene for FSCs (Horiguchi et al., 2010, Osuna et al., 2012). However, S100b is also expressed in MCL (Sasaki and Higuchi, 2022) and Sox2 has been shown to be expressed in parenchymal S100b-positive cells (Yoshida et al., 2011). In general, the findings that anterior pituitary stem cells express Sox2 and S100b are not sufficient to claim that all pituitary cells that express Sox2 are stem cells.

Our recent study suggests that parenchymal Sox2 cells, or a subset of these cells, are FSCs and that their differentiation is not associated with extensive reprogramming, unlike gonadotrophs. The study showed that *Sox2*-positive cells consist of two subclusters: a larger one, originally named FSC1, and a small one, originally named FSC2 (Fletcher et al., 2023). The basis for this designation was the co-expression of most genes in both FSC1 and FSC2, including *Aldh1a1, Aldh1a2, Aldh3a1, Capn6, Fmo1, Galm, Mt2A, Pde1c, Pla2g7, Slc6a8, S100b*, and *Timp1*. FSC2 also express genes not seen in FSC1, including *Cdh1, F3, Fosl1*, *Klf5, Krt17*, *Sfn*, and *Tent5b*. Additional data supported the hypothesis that the larger subcluster represents a parenchymal population of *Sox2*-expressing FSCs, and the smaller subcluster represents a population of *Sox2*-expressing stem cells in the marginal zone (Fletcher et al., 2023). These observations are consistent with the partial overlap of stem cells and FSCs, suggested by others (Vankelecom and Chen, 2014, Laporte et al., 2020). The heterogeneity of *Sox2*-expressing cells is also indicated by the presence of two clusters of these cells in the male mouse pituitary and the expression of several genes that we observed only in FSC2 (De Vriendt et al., 2025). Among MCL-specific genes, several lines of evidence suggest an important role that *Cdh1* encoding E-cadherin 1 may play in postnatal migration of Sox2/S100b cells in parenchyma (Fauquier et al., 2008, Lamouille et al., 2014, Batchuluun et al., 2017, Fujiwara et al., 2020).

Previous experiments have shown that PI4KA is critical for GnRH receptor signaling functions and hormone secretion of postnatal gonadotrophs, but it was unclear whether loss of this enzyme also affects gonadotroph differentiation from stem cells (Constantin et al., 2023). To address this question, we compared the transition of marginal cells to gonadotrophs in control and GSKO female mice. The data illustrate that the presence of gonadotrophs in MCL is significantly reduced in prepubertal GSKO mice, suggesting a role for PI4KA in postnatal gonadotrophs differentiation. This causes a decrease in gonadotroph density in the parenchyma that reflects the loss of tdTomato-positive cell in the pituitary by approximately 50%. This is a somewhat surprising finding, as the marginal cells are in good condition and initial gonadotroph differentiation should not be affected in GSKO pituitary, as embryonic gonadotroph differentiation is operative in these animals. Controlled death of somatotrophs leads to activation of stem cells and differentiation of somatotrophs (Fu et al., 2012). If target cells signal stem cells to increase its proliferation and differentiation into endocrine pituitary cells, we can assume that this signaling pathway is not operative in GSKO gonadotrophs.

In this study, we also observed two types of gonadotrophs: cells expressing tdTomato+LHB and cells expressing tdTomato alone. Both cell types were observed in controls and GSKO pituitaries. In the control group, the tdTomato+LHB subpopulation represented the majority of gonadotrophs, but progressively decreased in postnatal GSKO mice with development. In contrast, the number of cells expressing tdTomato alone was significantly higher in GSKO pituitary. We also observed a rapid loss of expression of the gonadotroph-specific genes *Gnrhr*, *Lhb* and *Fshb* in male and female GSKO pituitaries. Therefore, the increase in the percentage of gonadotrophs positive only for tdTomato in GSKO pituitaries reflects their regression due to the progressive loss of their differentiated state.

PI4KA knockout should affect the content of PI(4,5)P2 and PI(3,4,5)P3 in the plasma membrane (Stojilkovic and Balla, 2023). Here we show that postnatal differentiation of gonadotrophs from stem cells requires PI4KA. Others have reported that GnRH and gonadal hormones are not needed for this process (Sheridan et al., 2025). Therefore, a decrease in PI(3,4,5)P3 content, rather than PI(4,5)P2 content, could be responsible for the reduction in gonadotroph differentiation rate in the GSKO pituitary. Cell lifespan also depends on PI(3,4,5)P3 and its kinase signaling (Cariboni et al., 2015), the plasma membrane content of PI(3,4,5)P3 should be reduced in GSKO gonadotrophs.

Furthermore, GnRH application failed to trigger the calcium signaling pathway and accompanied electrical activity in GSKO gonadotrophs, and injection of InsP3 rescues this function in a fraction of cells (Constantin et al., 2023). This indicates that GSKO reduced plasma membrane PI(4,5)P2 below the level required for the phospholipase C-derived InsP3 threshold. This renders the GnRH receptor coupling to the Gq/11 signaling pathway ineffective, causing a progressive decrease in gonadotroph-specific gene expression, followed by loss of GnRH receptors and gonadotropin proteins. Reduced InsP3 production may affect the lifespan of tdTomato cells in GSKO mice, as has been reported for other PI(4,5)P2-depleted cell types (Mejillano et al., 2001).

## 5. Conclusion

In this study, we confirmed the existence of three populations of Sox2-positive cells in the mouse pituitary: parenchymal Sox2 single and small cluster cells, Sox2 cells constituting the MCL, and Sox2 expression by pituicytes. We visualized the functions of Sox2 cells in the MCL as stem cells for gonadotroph differentiation and their migration into the parenchyma. Gonadotroph differentiation was most pronounced during the juvenal-prepubertal period and was virtually silent in the adult pituitary. Juvenile - prepubertal gonadotroph differentiation was disrupted by PI4KA knockout in these cells, followed by a progressive loss of function of residual gonadotrophs, caused by the loss of gonadotroph-specific gene expression and protein synthesis. It is unlikely that parenchymal cells expressing Sox2 contribute to gonadotroph differentiation, and these cells or a subset of these cells appear to be differentiated FSCs. Further studies are needed to clarify whether/when Sox2 cells in MCL differentiate into other hormone-producing cells and/or FSCs and whether FSCs and pituicytes retain proliferative function.

## CRediT authorship contribution statement

**Kosara Smiljanic:** Writing – original draft, Writing - review & editing, Methodology, Investigation, Conceptualization. **Stephanie Constantin**: Writing – review & editing, Methodology, Investigation, Formal Analysis, Conceptualization. **Naseratun Nessa:** Writing – review & editing, Investigation. **Stanko S. Stojilkovic**: Writing – original draft, Writing - review & editing, Visualization, Supervision, Project administration, Conceptualization.

## Data availability

Data will be made available on request.

## Acknowledgements

This work was supported by National Institutes of Health grants from the Intramural Research Program of the *Eunice Kennedy Shriver* National Institute of Child Health and Human Development, award number: Z01 HD000195; grant recipient S.S.S. The floxed PI4KA mouse strain was provided by GlaxoSmithKline LLC under a Materials and Cooperative Research and Development Agreement. Confocal imaging was performed at the NICHD Microscopy and Imaging Core of the NIH, with the kind assistance of Drs. Vincent Schram and Ling Yi.

